# Age-specific pediatric templates improve EEG source-reconstructed biomarkers in children

**DOI:** 10.64898/2026.11.04.764182

**Authors:** Marit van den Berg, Tina Ganjipour, Marina Diachenko, Maria Candellero, Huibert D. Mansvelder, Hilgo Bruining, Arthur-Ervin Avramiea, Klaus Linkenkaer-Hansen

**Author notes:** Correspondence: Klaus Linkenkaer-Hansen. These authors contributed equally to this work (Marit van den Berg, Tina Ganjipour). These authors share senior authorship (Arthur-Ervin Avramiea, Klaus Linkenkaer-Hansen).

## Abstract

EEG source reconstruction requires an anatomical head model, ideally constructed from individual magnetic resonance imaging (MRI) scans. In developmental studies of children and adolescents, however, MRIs are rarely available. Researchers therefore routinely rely on adult templates, such as the FreeSurfer average (fsaverage), despite substantial anatomical changes throughout development. How much this reliance on adult models biases downstream EEG biomarkers remains unclear. Here, we investigated whether age-specific pediatric templates offer a better alternative to fsaverage for EEG-derived biomarkers in children and adolescents, using individual MRI-based reconstruction as the reference. We analyzed eyes-closed resting-state EEGs from 384 Healthy Brain Network participants aged 5.5 to 17.5 years. For each participant, we tested three anatomical models: their individual MRI, an age-specific pediatric template from a series spanning ages 6–17 at one-year intervals, and the adult fsaverage template. We compared four EEG biomarkers in childhood and adolescence: absolute power, relative power, long-range temporal correlations quantified with detrended fluctuation analysis (DFA), and functional excitation-inhibition (fE/I) ratio. Pediatric templates agreed more closely with individual MRI-based biomarkers in both developmental groups and had fewer parcels with significantly different biomarker values than the adult template, except for absolute power in older adolescents. We conclude that when individual MRI is not available, age-specific pediatric templates match individual MRI-based EEG biomarker values more closely than the adult template across all four biomarkers up to approximately 12 years of age and should be the preferred choice at least up to this age.

## 1 Introduction

Electroencephalography (EEG) is a non-invasive method for measuring and recording the brain’s electrical activity with high temporal resolution using electrodes attached to the scalp; however, identifying the anatomical origin of the activity is challenged by spatial blurring due to volume conduction and the superposition of signals from multiple neuronal populations (Burle et al., 2015). EEG source reconstruction can improve spatial interpretation by estimating the location of underlying brain signals (Michel et al., 2004). Head modeling is a crucial step in EEG source reconstruction because the brain, skull, and scalp conduct electricity differently and influence how electrical potentials reach the electrodes (Hallez et al., 2007). Inaccuracies in head modeling can lead to source localization errors (Acar & Makeig, 2013; Vorwerk et al., 2019; Montes-Restrepo et al., 2014), potentially affecting downstream EEG biomarker estimates. The most accurate head models are obtained using individual magnetic resonance imaging (MRI) (Acar & Makeig, 2013). However, these are rarely available, and anatomical templates are therefore often used.

A commonly used template is fsaverage, which represents the average adult brain and is incorporated into EEG source-reconstruction pipelines using FreeSurfer (Fischl, 2012) and MNE-Python (Gramfort et al., 2013, 2014). Fsaverage provides surfaces representing the boundaries between brain, skull, and scalp needed for head modeling, as well as cortical surfaces for defining source locations. MNE provides the corresponding precomputed BEM solutions. Since fsaverage represents the average adult brain, it does not accurately capture pediatric anatomy. Studies show that brain structure undergoes developmental changes throughout childhood and adolescence, including changes in cortical folding (Klein et al., 2014) and head and skull morphology (Gajawelli et al., 2020). Therefore, using an adult template for EEG source reconstruction in children and adolescents can introduce inaccuracies due to these developmental differences in brain anatomy and structure. Although several pediatric templates have been developed (Wilke et al., 2008; Fonov et al., 2011; Molfese et al., 2021; Richards et al., 2016; Sanchez et al., 2012), how template choice affects EEG source-reconstructed biomarker values in children and adolescents compared to individual MRI-based reconstruction remains unclear.

Here, we investigate how anatomical modeling choices affect EEG biomarker values derived from source reconstruction across childhood and adolescence. To address this aim, EEG source reconstruction was performed for each participant using three anatomical modeling strategies: an individual MRI-based head model, an age-specific pediatric template from the Neurodevelopmental MRI database (Richards et al., 2016), and the standard adult template from FreeSurfer, fsaverage. We used individual MRIs and EEG recordings from the open-source Healthy Brain Network database (Alexander et al., 2017). We compared EEG-derived biomarkers obtained with the pediatric and adult templates with those obtained using individual MRI-based reconstruction, which served as the reference. The biomarkers included the widely used absolute and relative power, as well as two brain dynamics algorithms: detrended fluctuation analysis (DFA) for quantifying long-range temporal correlations (Linkenkaer-Hansen et al., 2001; Hardstone et al., 2012) and excitation-inhibition balance assessed using the functional excitation-inhibition ratio (fE/I) (Bruining et al., 2020; Diachenko et al., 2024). These findings may help researchers choose a more appropriate anatomical template for developmental groups when individual MRI is unavailable.

## 2 Methods

### 2.1 Data sources

#### 2.1.1 HBN

For this study, we analyzed T1-weighted MRI, EEG, and phenotypic data from the Healthy Brain Network (HBN), a pediatric biobank (ages 5–21) including both typically developing participants (“healthy”) and individuals with atypical neurodevelopmental profiles (Alexander et al., 2017). The phenotypic data were accessed under the HBN Data Usage Agreement. The detailed inclusion and exclusion criteria as well as the protocols can be found in the original HBN paper (Alexander et al., 2017). The original HBN study was approved by the Chesapeake Institutional Review Board. Written informed consent was obtained from parents or legal guardians for participants younger than 18 years, with written assent obtained from the participants (Alexander et al., 2017). The present study involved secondary analysis of previously collected data, and no additional data were collected.

We included participants when a high-quality T1-weighted MRI scan, a cleaned resting-state EEG recording, and phenotypic data were available. Additionally, only participants between the ages 5.5 and 17.5 were included. A total of 414 participants met the initial inclusion criteria; 30 participants were then excluded during head model construction (16 out of 183 in childhood and 14 out of 231 in adolescence). The final analysis cohort comprised 384 participants with successful source reconstruction for all three anatomical modeling strategies (Table 1). The age and sex distributions of the final cohort are shown in Supplementary Figure S1.

**Table 1.** Demographics of the HBN cohort.

| **Characteristic** | **Childhood** | **Adolescence** | **Total** |
|---|---|---|---|
| Number of participants, *n* | 167 | 217 | 384 |
| Age range (years) | 5.5– <10.5 | 10.5–17.5 | 5.5–17.5 |
| Age, mean ± SD (years) | 8.49 ± 1.28 | 13.79 ± 2.01 | 11.49 ± 3.15 |
| Male, *n* (%) | 106 (63.5%) | 151 (69.6%) | 257 (66.9%) |
| Female, *n* (%) | 58 (34.7%) | 66 (30.4%) | 124 (32.3%) |
| Other, *n* (%) | 3 (1.8%) | 0 (0.0%) | 3 (0.8%) |
| Healthy, *n* (%) | 13 (7.8%) | 22 (10.1%) | 35 (9.1%) |

#### 2.1.2 Neurodevelopmental MRI database template

We used age-specific pediatric MRI templates spanning 6 to 17 years at one-year intervals from the John E. Richards Lab’s Neurodevelopmental MRI Database (Evans, 2006; Sanchez et al., 2010, 2012; Richards & Xie, 2015; Richards et al., 2016) to create age-specific head models for EEG source reconstruction. Participants were assigned to the pediatric template corresponding to their age rounded to the nearest whole year. Detailed inclusion and exclusion criteria as well as the protocols can be found in Evans (2006) and Richards et al. (2016).

### 2.2 MRI acquisition HBN

For this study, T1-weighted structural MRIs from the HBN (releases 1–9) were obtained using DataLad (version 1.3.1) (Halchenko et al., 2021) from the Reproducible Brain Chart GitHub repository named *HBN_BIDS* (https://github.com/ReproBrainChart/HBN_BIDS). All images had previously undergone quality control using the MRI Quality Control Tool (Esteban et al., 2017). MRI scans were marked as either “Pass”, “Artifact”, or “Fail”. For this study, we only included scans that were marked as “Pass” to minimize the influence of imaging artifacts.

Participants underwent MRI scanning at one of four sites: Staten Island, using a mobile 1.5 T Siemens Avanto; Rutgers University Brain Imaging Center, using a 3.0 T Siemens Tim Trio; or Citigroup Cornell Brain Imaging Center and CUNY Advanced Science Research Center, both using a 3.0 T Siemens Prisma.

### 2.3 EEG data acquisition and processing HBN

EEG data were recorded using a 128-channel EEG Geodesic HydroCel system (Electrical Geodesics Inc.) at a sampling rate of 500 Hz with a 0.1 to 100 Hz bandpass (Alexander et al., 2017). Participants alternated between eyes closed and eyes open. Procedural details can be found in the original HBN paper (Alexander et al., 2017). The EEG data were cleaned for artifacts by trained students at the CNCR-INF department in Amsterdam using Robin’s Viewer (Weiler et al., 2023). This included an initial visual inspection and removal of high-amplitude artifactual segments, identification and interpolation of bad channels, average re-referencing, Independent Component Analysis (ICA) to remove repetitive artifacts (e.g., eye blinks, lateral eye movements, heartbeat, and sustained muscle activity), followed by a final visual inspection to remove any residual artifactual fragments. Only the eyes-closed resting-state condition was included in downstream biomarker analyses to minimize the influence of visual processing on EEG activity. An overview of the complete study workflow is shown in Figure 1.

**Figure 1.**
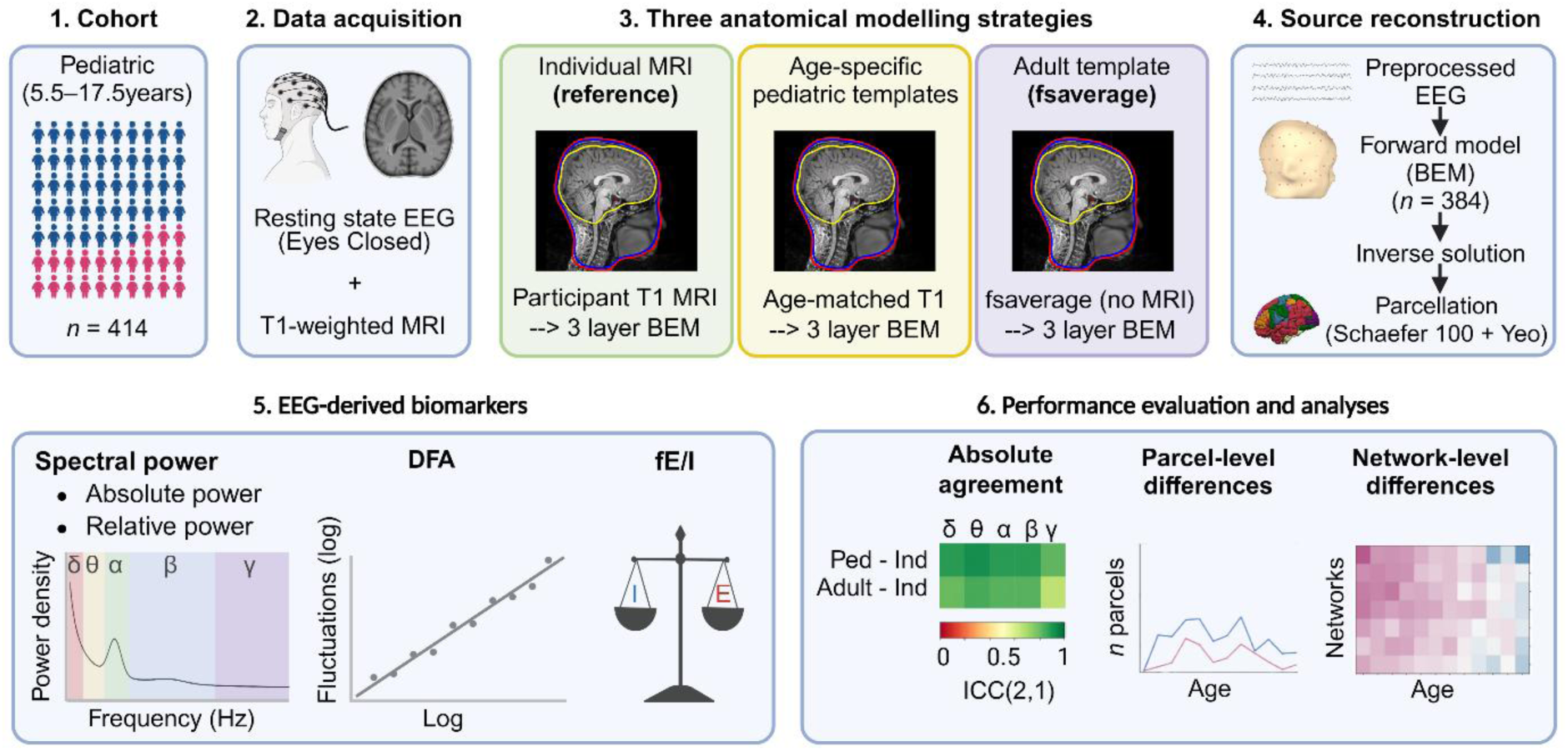
Study workflow. Resting-state EEG data from 414 participants were used for source reconstruction using three anatomical modeling strategies: individual MRI-based head models derived from T1-weighted MRI, age-specific pediatric template-based head models derived from the Neurodevelopmental MRI Database, and the adult template-based head models (fsaverage). Following source reconstruction, EEG-derived biomarkers (spectral power, detrended fluctuation analysis, and functional excitation-inhibition ratio) were computed and compared for the final cohort of 384 participants across the three modeling strategies using absolute agreement, parcel-level paired *t*-tests, and network-level absolute mean deviations. Created in BioRender. Van Den Berg, M. (2027) https://BioRender.com/ezlik4u.

### 2.4 Head model construction

#### 2.4.1 Individual MRI model (reference)

The structural T1-weighted MRIs were processed using the FreeSurfer recon-all pipeline (version 8.1.0) (Fischl, 2012) to reconstruct individual cortical surfaces and create anatomical segmentations needed for EEG source reconstruction. To construct a three-layer boundary element model (BEM) of the head, we used MNE’s watershed-BEM algorithm (v1.12.1) to generate the inner skull, outer skull, and outer skin surfaces (Gramfort et al., 2013, 2014). To improve surface generation, a watershed preflood height of 20 (*h* = 20) and a brain surface shrink factor of 3 (*shk_br_surf* = 3) were used during watershed segmentation. If intersections occurred between the inner and outer skull, or outer skull and outer skin, then these were corrected using an approach adapted from the *correct_intersecting_meshes* function in the code from O’Reilly et al. (2021). This approach repositioned vertices of the outer surface in each pair, using a correction margin of 0.5 mm. Among the included participants, 56 out of 167 (33.5%) for childhood and 17 out of 217 (7.8%) for adolescence required BEM surface corrections. The three-layer BEM incorporated ico-4 surface tessellation and default conductivity values (= 0.3, 0.006, 0.3 S/m). Identical conductivity values were used across models to isolate the effect of anatomical modeling choice. A cortical source space was generated using ico-5 surface tessellation.

Quality control was performed at several stages of head model construction. First, FreeSurfer processing logs were reviewed to identify incomplete runs. Second, the three BEM surfaces generated using the watershed algorithm (inner skull, outer skull, and outer skin) were visually evaluated in Freeview by overlaying the surfaces on the subject’s T1 MRI. Completion of BEM model construction and BEM solution computation was assessed using automated scripts that checked the processing logs for success messages, additionally checking for the failure message indicating that the inner skull surface was not completely inside the outer skull surface. Corrected surfaces were reassessed using visual inspection and automated checks. Examples of corrected surfaces and surfaces leading to exclusion are shown in Supplementary Figure S2. Source space generation was assessed using an automated script to check the logs for success messages and a visual inspection using MNE’s visualization tools. Lastly, the coregistration between EEG and MRI was visually evaluated using MNE’s 3D coregistration interface to inspect the alignment between electrode positions and the scalp surface. Participants whose head models failed quality control after any corrections were excluded from further analyses. Incomplete FreeSurfer runs only led to exclusion when failures affected anatomical structures required for subsequent source-reconstruction steps.

#### 2.4.2 Adult template model (fsaverage)

The fsaverage template, a precomputed adult anatomical model provided by FreeSurfer, served as the adult template (Fischl, 2012; Fischl et al., 1999). All other source-reconstruction parameters were kept the same as described in the individual MRI pipeline, including default conductivity values, ico-4 BEM, and ico-5 cortical source space.

#### 2.4.3 Pediatric template model

For the pediatric template-based models, we used age-specific anatomical templates from the Neurodevelopmental MRI Database. We extracted the three BEM surfaces (inner skull, outer skull, and outer skin) from the template BEM segmentation volume files following the approach of O’Reilly et al. (2021). The three labels were combined into nested masks representing the inner skull, outer skull, and outer skin surfaces. MNE’s marching cubes algorithm generated the surface meshes. The resulting surfaces were then transformed to FreeSurfer RAS coordinates, smoothed, and decimated. If intersections occurred between the pial and inner skull, or inner and outer skull, or outer skull and outer skin, they were corrected as described in Section 2.4.1. A three-layer BEM model was subsequently constructed using conductivity values of 0.3, 0.006, and 0.3 S/m for scalp, skull, and brain, respectively, using MNE with *ico* = None. Cortical surfaces provided with each template were used to generate an ico-5 source space.

### 2.5 Forward and inverse modeling

Digitized electrode positions were unavailable, so we used standard HydroCel montage positions for all participants and excluded channels E48, E119, E126, and E127 from source reconstruction. Coregistration aligned EEG electrode locations with each anatomical model, using participant-specific transformations for individual MRIs and template-specific transformations for the adult and pediatric templates. For all three models, an initial automatic fiducial fit was followed by refinement using the iterative closest point (ICP) algorithm, with a maximum of 100 iterations, with EEG weight set to 1, and all other weights set to 0. The rigid transformations involved no scaling of the anatomical models. All coregistration results were visually inspected.

The forward solution was computed using *mne.make_forward_solution* (Gramfort et al., 2013, 2014), incorporating the BEM, source space, coregistration, EEG electrode locations, and channel information. A minimum distance of 5.0 mm was used to exclude source points (dipoles) located too close to the inner skull surface.

A minimum norm inverse solution provided estimates of source activity. The inverse operator was computed with an ad hoc noise covariance using the default EEG value of 0.2 µV, free orientation (*loose* = 1), and the default depth parameter of 0.8. The inverse solution used a signal-to-noise ratio (SNR) of 3, and a λ² of 1/9, and vector orientation.

### 2.6 Parcellation: Schaefer and Yeo atlas

We parcellated the cortex into 100 parcels using the Schaefer 2018 atlas (Schaefer et al., 2018). For the individual MRI-based models, the atlas labels were morphed from fsaverage to each subject. The adult template-based model used the atlas labels directly, whereas the pediatric template-based models required morphing the labels from fsaverage to each pediatric template. We computed a mean time series for each parcel using sign-flip averaging (*mean_flip*). The Schaefer atlas labels corresponded to seven functional networks defined by Yeo et al. (2011) in each hemisphere: Visual, Somatomotor, Default, Limbic, Dorsal Attention, Ventral Attention, Frontoparietal.

### 2.7 Biomarker computation

We computed EEG biomarkers for each parcel and frequency band. The five classical frequency bands spanned 1–45 Hz: delta (1–4 Hz), theta (4–8 Hz), alpha (8–13 Hz), beta (13– 30 Hz), and gamma (30–45 Hz). Biomarkers included absolute power, relative power, detrended fluctuation analysis (DFA), and the functional excitation-inhibition ratio (fE/I).

#### 2.7.1 Spectral power

We computed the power spectral density (PSD) over 1–45 Hz using Welch’s method with a Hamming window, implemented in the MNE software (Gramfort et al., 2013, 2014). The length of the Fast Fourier Transform (FFT) was set to ten times the sampling frequency, resulting in a frequency resolution of 0.1 Hz. Each segment used a Hamming window equal in length to the FFT length, and window overlap was set to 50% of the FFT length, rounded down to the nearest integer. We calculated absolute power for each of the five frequency bands by integrating the PSD over the respective band and converting the result to a decibel (dB) scale. Relative power was computed as the ratio of each band’s integrated PSD to the total integrated PSD across the full 1–45 Hz frequency range and expressed in %.

#### 2.7.2 Detrended fluctuation analysis (DFA)

We quantified autocorrelation in the amplitude envelope of each frequency band using the DFA exponent, which relates mean amplitude fluctuations to increasing time-window sizes within a chosen time-scale range (Linkenkaer-Hansen et al., 2001; Hardstone et al., 2012). A DFA exponent of 0.5 indicates an uncorrelated random signal, whereas values >0.5 denote positive autocorrelation whose strength grows with the exponent. Before computing DFA, each signal was bandpass filtered into five frequency bands, and the amplitude envelope for each band was extracted using a Hilbert transform. Bandpass filtering and Hilbert transformation were implemented in the MNE software (Gramfort et al., 2013, 2014). For each frequency band, the DFA exponent was obtained as the slope of the line relating the log-fluctuation to the log-window size, fitted using the time-scale ranges optimized by Diachenko et al. (2024). A step-by-step description of this algorithm is provided in Supplementary Material S1 of Diachenko et al. (2024).

#### 2.7.3 Functional excitation-inhibition ratio (fE/I)

We used fE/I, which captures the covariance between the signal’s amplitude and its autocorrelation, to quantify the network-level balance between excitation and inhibition (Bruining et al., 2020). The fE/I value <1 indicates inhibitory dominance in the signal, >1 excitatory dominance, and 1 a balanced E/I ratio. Before computing fE/I, each signal was bandpass filtered into five frequency bands, and the amplitude envelope for each band was extracted using a Hilbert transform (also via the MNE software). For each band, the temporal variation in amplitude autocorrelation was approximated by windowed amplitude fluctuations, and fE/I was calculated as 1 minus the Pearson correlation between the windowed amplitudes and their fluctuations. The DFA exponent was computed as described above, and fE/I was set to missing (NaN) whenever DFA <0.6, because the covariance estimate is only meaningful for signals with substantial autocorrelation (Diachenko et al., 2024; Bruining et al., 2020). A step-by-step description of this algorithm is provided in Supplementary Material S2.1 of Diachenko et al. (2024).

### 2.8 Evaluation in the HBN cohort

For each participant, we performed source reconstruction using three head modeling strategies: an individual MRI-based model, an adult template-based model (fsaverage) and a pediatric template-based model. The individual MRI-based model served as the reference for evaluating both template-based models.

#### 2.8.1. EEG biomarker absolute agreement

We assessed agreement between template-based and individual MRI-based source reconstruction for absolute power, relative power, DFA, and fE/I. Using parcel-level biomarker values, we performed analyses separately for childhood (5.5 to <10.5 years) and adolescence (10.5 to 17.5 years), comparing adult template-based and pediatric template-based reconstructed biomarkers with individual MRI-based biomarkers.

We quantified absolute agreement using the intraclass correlation coefficient, ICC(2,1) (Shrout & Fleiss, 1979; McGraw & Wong, 1996; Koo & Li, 2016). For each biomarker and frequency band, ICCs were calculated across participants for each parcel and summarized using the median across parcels. Higher ICC values indicate greater absolute agreement between the models. The ICC was calculated as follows:

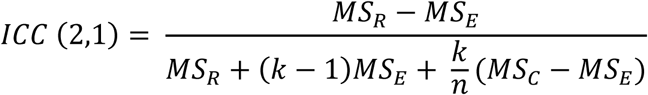

where MS_R_ is the mean square for rows (participants), MS_E_ is the residual mean square error, MS_C_ is the mean square between reconstruction methods, *n* is the number of participants, and *k* is the number of measurements being compared (here, *k* = 2). We calculated differences in agreement between comparisons as ΔICC = median ICC(Pediatric–Individual) − median ICC(Adult–Individual). Positive values indicate greater agreement for the pediatric templates. The 95% bootstrap confidence intervals were calculated using 10,000 paired participant-level resamples within each developmental group, recalculating parcel-wise ICCs, their medians across parcels, and ΔICC for each resample. Two-sided bootstrap *p*-values were adjusted using the Benjamini-Hochberg false discovery rate (FDR) correction, with an FDR threshold of *q* = 0.05, across 40 tests (2 developmental groups × 4 biomarkers × 5 frequency bands) (Benjamini & Hochberg, 1995). Adjusted *p*-values below 0.05 were considered significant.

#### 2.8.2. EEG biomarker parcel-level differences

To evaluate parcel-level differences in template performance across ages, we compared template-based biomarker values with individual MRI-based biomarker values using two-sided paired *t*-tests for each parcel, biomarker, frequency band, and one-year age bin. For each test, we included only participants with valid values for both reconstruction methods being compared. *P*-values were adjusted using the Benjamini-Hochberg FDR correction (*q* = 0.05) across parcels, separately for each biomarker, frequency band, age bin, and reconstruction method comparison. Adjusted *p*-values below 0.05 were considered significant. The number of participants contributing to the fE/I paired tests varied because fE/I values were only calculated when DFA ≥ 0.6. The participant counts are provided in Supplementary Table S1.

#### 2.8.3. Network-level deviations in absolute power

To evaluate network-level differences in template performance across ages, we calculated absolute deviations between template-based and individual MRI-based absolute power values for each participant, parcel, and frequency band. We then averaged these deviations across participants within one-year age bins, across the five frequency bands, and across parcels within each Yeo network. The difference was calculated as adult minus pediatric mean absolute deviation from individual MRI-based values. Positive values indicate that pediatric template-based values were closer to individual MRI-based values, whereas negative values indicate that adult template-based values were closer.

## 3 Results

In this study, we compared EEG-derived biomarkers obtained using three anatomical modeling strategies for each participant: an adult template, an age-specific pediatric template, and an individual MRI-based model. The cohort comprised 384 children and adolescents. We assessed absolute agreement, significant parcel-level differences, and network-level mean absolute deviations.

### 3.1 Pediatric templates show greater agreement with individual MRI-based biomarkers

To assess whether pediatric or adult template-based reconstructions showed greater agreement with individual MRI-based reconstructions at the biomarker level, we calculated absolute agreement intraclass correlation coefficients (ICCs) for absolute power, relative power, DFA and fE/I across five conventional frequency bands, separately for childhood (6– 10 years) and adolescence (11–17 years). We calculated ICCs per parcel and summarized them across parcels using the median (Fig. 2A, B). Examples of poor, moderate, and good agreement illustrate that median ICCs are representative of the parcel-wise ICC distributions (Fig. 2C–E).

**Figure 2.**
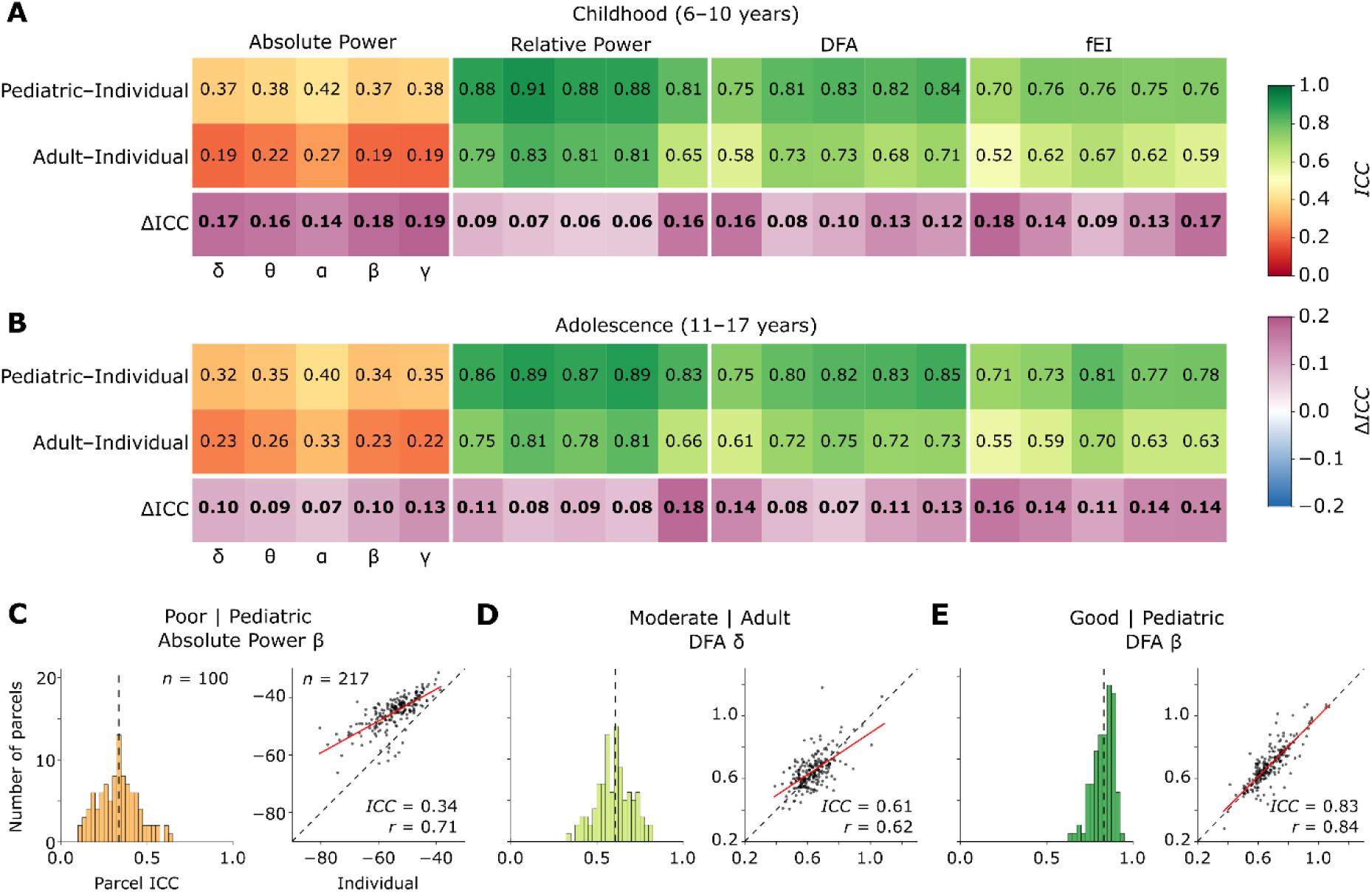
Pediatric templates show greater agreement with individual MRI-based EEG biomarkers than the adult template across developmental groups. (A,B) Absolute-agreement intraclass correlation coefficients (ICC(2,1)) were calculated per Schaefer parcel and summarized across parcels using the median for childhood (6 to 10 years; *n* = 167) and adolescence (11 to 17 years; *n* = 217). Higher ICC values indicate greater agreement with individual MRI-based reconstruction. The third row shows ΔICC = ICC(Pediatric,Individual) − ICC(Adult,Individual), with positive values (*purple*) indicating greater agreement for the pediatric templates. All ΔICC comparisons were significant after Benjamini-Hochberg FDR correction (*q* = 0.05). Biomarkers: absolute power, relative power, detrended fluctuation analysis (DFA), and functional excitation-inhibition ratio (fE/I). Frequency bands: δ = 1–4 Hz, θ = 4–8 Hz, α = 8–13 Hz, β = 13–30 Hz, and γ = 30–45 Hz. (C–E) Examples of poor (<0.50), moderate (0.50–0.75), and good (0.75–0.90) agreement between EEG biomarkers derived from the template and individual MRI reconstructions in the adolescent group. Panels C and E use the pediatric template; panel D uses the adult template. Histograms show ICC values across 100 parcels; the vertical *dashed line* marks the median in each plot. Scatterplots show individual MRI-based values on the x-axis and template-based values on the y-axis. Each point represents one participant (*n* = 217). *Red lines* show the least-squares fits, and the diagonal *dashed lines* show exact agreement (*y* = *x*). Panel C shows that Pearson correlation (r = 0.71) can exceed the absolute-agreement ICC (0.34).

Relative power, DFA, and fE/I showed moderate to good agreement in both children and adolescents, with the highest median ICCs observed for relative power when using the pediatric template (Fig. 2A,B). In contrast, absolute power showed poor agreement (median ICCs < 0.50) in both developmental groups, particularly when using the adult template. Overall, median ICCs between the pediatric template and individual MRI reconstructions were higher than those between the adult template and individual MRI reconstructions across all biomarkers and frequency bands in both developmental groups (Fig. 2A,B). These differences were significant for all biomarkers and frequency bands (Fig. 2A,B; ΔICC, all *p* < 0.05, FDR-corrected, *q* = 0.05). This indicates that pediatric templates provide EEG biomarker estimates that are more consistent with those obtained using individual MRI reconstructions than adult templates. Bootstrap 95% confidence intervals for the ICC differences are shown in Supplementary Figure S3.

The importance of assessing absolute agreement rather than correlation alone is further illustrated in Figure 2C. Although absolute power values derived from the pediatric template and individual MRI reconstructions showed a relatively high Pearson correlation (*r* = 0.71), their deviation from the identity line (*x* = *y*) revealed poor absolute agreement (ICC = 0.34).

### 3.2 Pediatric templates show fewer parcel-level biomarker differences during childhood

The previous analysis showed that pediatric templates provided greater agreement with individual MRI reconstructions than adult templates based on ICCs. To further investigate how template choice affects detectable biomarker differences, we next used paired *t*-tests to identify parcels in which template-based biomarker values differed significantly from those obtained using individual MRI reconstructions, separately for each age, biomarker, and frequency band.

In children, age-specific pediatric templates showed fewer parcel-level differences from individual MRIs than the adult template did across all four biomarkers (Fig. 3A–D). Absolute power showed the largest number of parcels with significantly different biomarker values, followed by relative power, whereas DFA and fE/I had the fewest. For fE/I, the number of participants included in the analysis varied per parcel, frequency band, age, and template (Supplementary Table S1).

**Figure 3.**
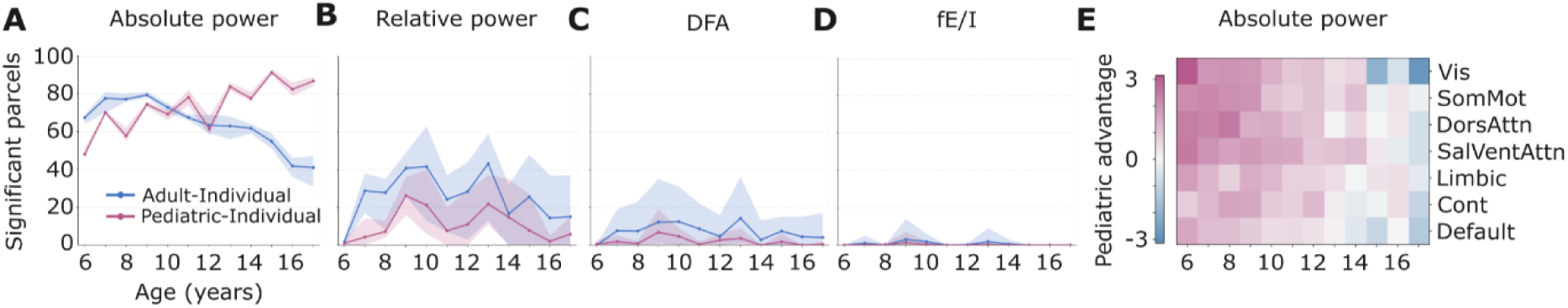
Pediatric templates show fewer biomarker differences than the adult template during childhood. **(A–D)** Number of Schaefer parcels showing significant differences from individual MRI-based reconstruction for absolute power (A), relative power (B), detrended fluctuation analysis (DFA; C), and functional excitation-inhibition ratio (fE/I; D) across age. Adult and pediatric templates were each compared with individual MRI-based reconstruction using paired *t*-tests with Benjamini-Hochberg FDR correction (*q* = 0.05) across 100 parcels per age, biomarker and frequency band. Lines show the mean across the five frequency bands, and the shaded areas show the minimum and maximum values. For absolute power, the adult template shows fewer parcels with significantly different biomarker values at older ages. **(E)** Adult minus pediatric mean absolute deviation from individual MRI-based reconstruction for absolute power across age and the seven Yeo networks. Deviations were averaged across participants, frequency bands, and parcels within each network. Positive values (*purple*) indicate that the pediatric template is closer to individual MRI-based reconstruction, whereas negative values (*blue*) indicate that the adult template is closer.

Overall, these results suggest that age-specific pediatric templates produce fewer parcel-level EEG biomarker differences than the adult template when compared to individual MRI-based reconstruction during childhood.

### 3.3 Template-related differences in absolute power vary with age

For absolute power, pediatric templates showed fewer parcels with significantly different biomarker values than the adult template during childhood. During adolescence, this number increased for the pediatric-individual MRI comparison but decreased for the adult-individual MRI comparison (Figure 3A). To examine whether this pattern in absolute power during adolescence was also reflected in the magnitude and location of absolute-power differences, we compared the mean absolute deviation of each template’s absolute-power values from individual MRI-based values across cortical networks.

Figure 3E shows that the pediatric template had a smaller mean absolute deviation from individual MRI-based absolute power at younger ages. This advantage of the pediatric template started to diminish around age 14. The strongest shift towards the adult template occurred at age 17 in the visual network.

## 4 Discussion

We investigated how the choice of anatomical model affects EEG source-reconstructed biomarkers across childhood and adolescence by comparing biomarker values obtained using pediatric and adult templates with those derived from individual MRI-based reconstructions. Age-specific pediatric templates showed greater absolute agreement with individual MRI-based biomarker values than the adult template across developmental groups. However, absolute agreement varied by biomarker. Absolute power showed poor absolute agreement (median ICC < 0.50) across all frequency bands and developmental groups, whereas relative power, DFA, and fE/I showed moderate-to-good agreement. In children, pediatric templates also generally showed fewer parcels with significant differences from individual MRI-based reconstructions than the adult template across all biomarkers. This pattern persisted into adolescence for relative power, DFA, and fE/I but reversed for absolute power. From approximately 14 years of age onward, the adult template showed fewer parcels with significant absolute-power differences relative to individual MRI-based reconstructions. Mean absolute deviations in absolute power across cortical networks showed a similar shift, with the adult template showing its strongest advantage at age 17 in the visual network.

The pediatric template advantage aligns with the idea that adult templates do not capture developmental changes in brain anatomy, such as changes in cortical folding (Klein et al., 2014) and head and skull morphology (Gajawelli et al., 2020), whereas age-specific pediatric templates should better reflect anatomy during development. A previous study in infants found that age-specific templates preserved the topographical distribution of source activity obtained using individual MRI, but failed to reproduce significant stimulus effects (Fu & Richards, 2022). They reported that age-inappropriate templates preserved neither and showed the lowest correlations with estimates from individual MRI. Our findings in children and adolescents similarly support the use of age-specific templates and suggest that their performance depends on the outcome assessed, with agreement varying across biomarkers. O’Reilly et al. (2021) similarly found that sources reconstructed using an adult template differed substantially from sources reconstructed using infant templates. Our findings extend this evidence for the importance of age-specific templates to resting-state EEG biomarker values in childhood and adolescence.

Absolute power showed the poorest agreement, suggesting that the amplitude of the reconstructed signal may be sensitive to the choice of anatomical model. Previous research has shown that different head models that include or exclude particular tissue compartments can produce different reconstructed source amplitudes in the same brain regions (Conte & Richards, 2021). Similarly, Ignatiadis et al. (2022) showed that using individual anatomy rather than template anatomy affected the amplitude of reconstructed auditory evoked responses. Together, these findings are consistent with our observation that absolute power is sensitive to anatomical modeling choice.

Although ICCs for the adolescent group favored the pediatric templates, at older ages the adult template yielded smaller mean absolute deviations from the individual MRI-based results for absolute power. The decrease in the pediatric advantage for absolute power with increasing age may reflect the anatomy becoming more similar to adult anatomy, consistent with the decrease in parcels with significantly different biomarker values between adult and individual MRI-based reconstruction (Fig. 3A). However, this does not explain the increase in parcels with significantly different biomarker values between pediatric and individual MRI-based reconstruction. One possible contributor is poorer EEG-MRI coregistration at older ages, with larger distances between the template electrode positions and the pediatric template scalp surfaces (Supplementary Table S2). Coregistration errors can affect source localization. Acar and Makeig (2013) demonstrated that a 5° tilted electrode montage can produce up to 12 mm localization errors. Digitized electrode positions can reduce source localization discrepancies (Liu et al., 2023). These findings concern localization accuracy; whether coregistration errors also explain the absolute-power deviations observed here remains unclear. Digitized electrode positions were not available in our study, so coregistration relied on a template montage.

The relative robustness of DFA and fE/I to changes in head models (Fig. 3C,D) is notable, although not entirely surprising. Reconstructing cortical currents at slightly different orientations or depths may affect the overall strength of the estimated source activity, as reflected in absolute power. However, the modulation of signal amplitude may be less affected, and this is an important property of the signal quantified by the DFA and fE/I algorithms (Linkenkaer-Hansen et al., 2007).

A strength of our study is that we compared all three modeling strategies within each participant, using the same EEG recording and constructing head models with methods that were as consistent as possible. Nevertheless, several factors can introduce variability into the individual MRI-based reconstructions used as our reference. First, head motion during MRI acquisition can bias anatomical estimates, including underestimation of gray matter volume and cortical thickness (Reuter et al., 2015). Differences in image quality may affect surface-based reconstruction even among scans rated as good quality (White et al., 2018). Furthermore, Mitrovska & Haufe (2025) showed that defacing of MRI scans, the removal of facial features to protect privacy, may affect EEG source localization. In our dataset, defacing was performed before we received the data. We also used the same fixed conductivity values across pipelines to maintain consistency. However, this does not account for any individual differences in skull conductivity (Antonakakis et al., 2020). Additionally, BEM surfaces required correction more frequently in younger participants, and some surface irregularities remained after correction despite passing the quality checks. We did not assess the effects of these factors independently.

Individual MRI-based reconstruction served as an anatomical reference rather than ground truth. Depuydt et al. (2024) used simulations to quantify localization error using template-based and individual MRI-based head models. A similar approach could be used in future work to extend our assessment of agreement by using simulations with known source activity to evaluate the accuracy of reconstructed biomarker values. Our findings suggest that when an individual MRI is not available, age-specific pediatric templates could be a suitable alternative to adult templates for EEG source-reconstructed biomarker analyses in pediatric cohorts. Specifically, these findings support their use for relative power, DFA, and fE/I across developmental groups, and for absolute power during childhood.

## Declaration of the use of AI

AI was used to check spelling and grammar. We used AI to reword our draft text.

## Data and Code Availability

The EEG and T1-weighted MRI data used in this study are publicly available through the Healthy Brain Network database. Phenotypic data can be accessed through the HBN-dedicated instance of LORIS or the COINS Data Exchange. The age-specific pediatric templates were constructed using anatomical resources obtained from the Neurodevelopmental MRI

Database (Richards et al., 2016), which is available through NITRC after registration and membership to the group. Code used for head model construction and statistical analyses is available from the corresponding author upon reasonable request and will be made publicly available in a repository upon acceptance of the manuscript.

## Author contributions

Conceptualization: KLH, MD

Methodology: MB, TG, KLH, MD, AEA

Formal analysis: MB, TG

Software: MB, TG, AEA

Data curation: MB, TG, MD, MC, AEA

Visualization: MB, TG, KLH, MD, MC

Project administration: KLH

Funding acquisition: KLH

Supervision: KLH, MD

Writing—original draft: MB, TG

Writing—review & editing: KLH, MD, AEA, MC, HB, HDM

## Acknowledgements

We thank the participants and their families, as well as the teams who collected and shared the data through the Healthy Brain Network and the Neurodevelopmental MRI database. We thank the NOC group, particularly the students involved in EEG data cleaning.

## Supplementary material

Supplementary material related to this article can be found in the online version.

## Supplementary materials

**Supplementary Figure S1.**
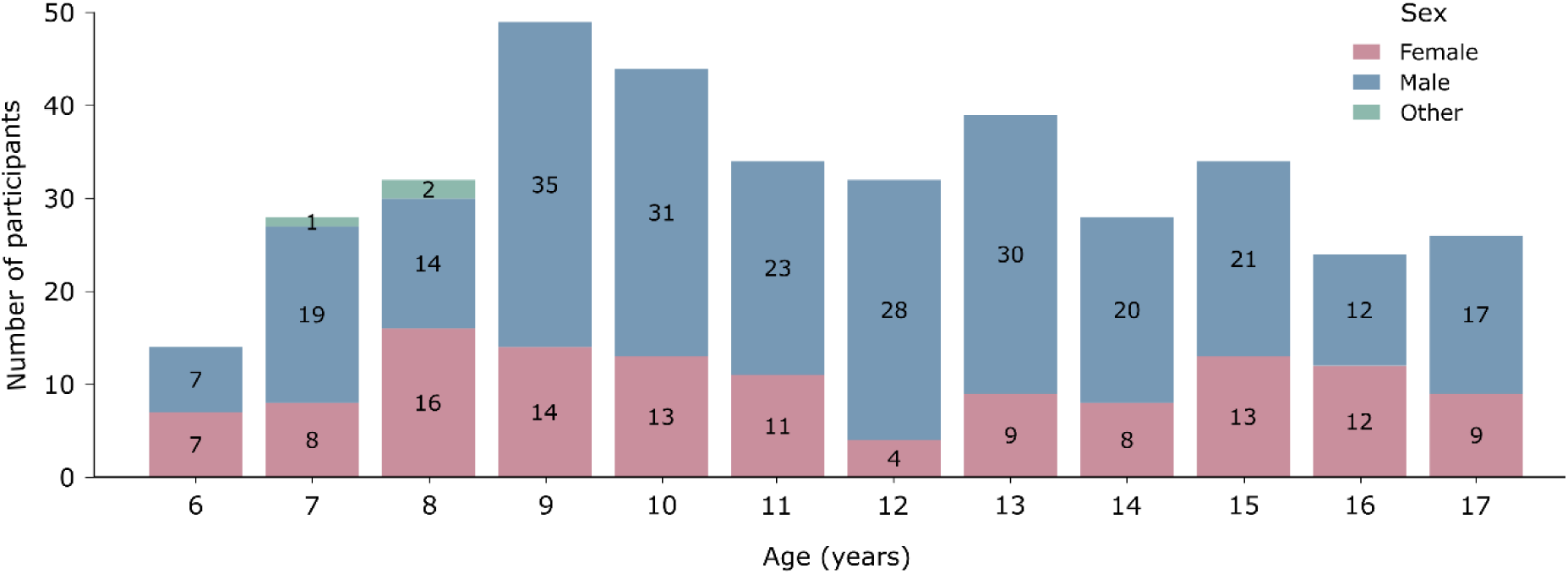
Age distribution by sex of participants included in the study. Number of participants per one-year age bin. Female participants are shown in pink, male participants in blue, and participants categorized as other in green. Numbers within the bars indicate participant counts.

**Supplementary Figure S2.**
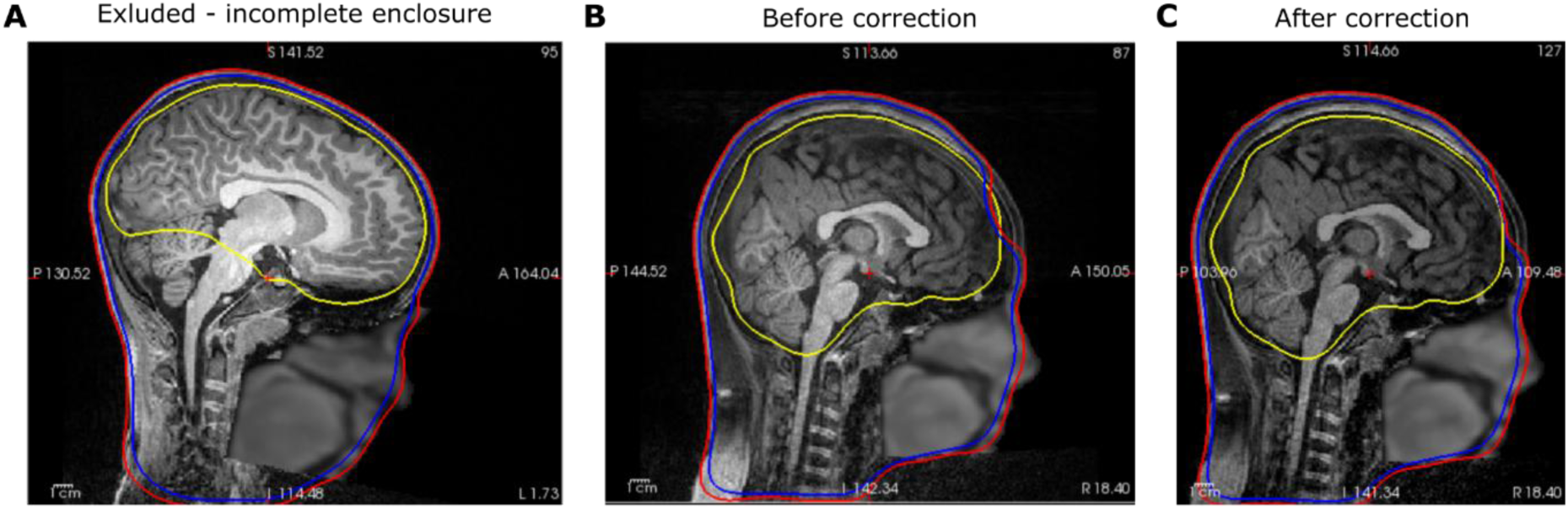
Examples of BEM exclusion and correction. Boundary Element Model (BEM) surfaces are shown overlaid on their corresponding T1-weighted MRI in sagittal view. Inner skull *(yellow line)*, outer skull (*blue line),* and outer skin surface (*red line)*. **(A)** Example leading to exclusion because the inner skull surface failed to enclose the entire brain, leaving out the cerebellum and brainstem. **(B, C)** Example of a participant whose inner skull surface was not fully contained within the outer skull surface, shown before (B) and after (C) intersection correction.

**Supplementary Figure S3.**
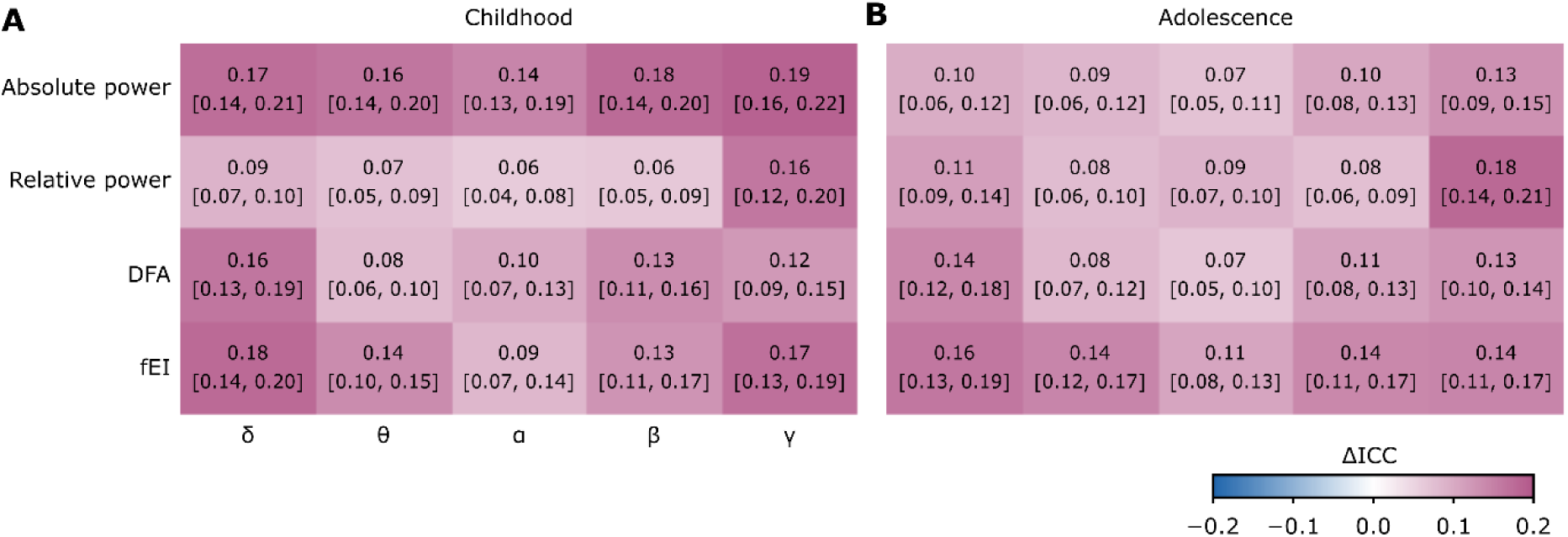
Bootstrap 95% confidence intervals for differences in ICC between pediatric and adult template comparisons. **(A, B)** Differences in intraclass correlation coefficients (ΔICC) were calculated as ICC(Pediatric–Individual) minus ICC(Adult– Individual). Positive values (*purple*) indicate greater agreement for the pediatric template. Values in each cell show ΔICC; brackets show their 95% bootstrap confidence intervals for childhood (A) and adolescence (B) across the four biomarkers: absolute power, relative power, detrended fluctuation analysis (DFA), functional excitation-inhibition ratio (fE/I) and five frequency bands. Frequency bands: δ = 1–4 Hz, θ = 4–8 Hz, α = 8–13 Hz, β = 13–30 Hz, and γ = 30–45 Hz.

**Supplementary Table S1.** Number of participants contributing to paired t-tests of fE/I parcel values.

| **Age (years)** | **Participants in age group, N** | **Adult-Individual median *n* (range)** | **Pediatric-Individual median *n* (range)** |
|---|---|---|---|
| 6 | 14 | 9 (3–14) | 9 (3–14) |
| 7 | 28 | 18 (4–28) | 19 (6–28) |
| 8 | 32 | 20 (9–32) | 21 (10–32) |
| 9 | 49 | 32 (11–48) | 33 (12–48) |
| 10 | 44 | 30 (11–43) | 32 (16–44) |
| 11 | 34 | 23 (8–34) | 25 (12–34) |
| 12 | 32 | 21 (7–32) | 23 (9–32) |
| 13 | 39 | 25 (9–39) | 26.5 (11–39) |
| 14 | 28 | 18 (7–27) | 19 (8–27) |
| 15 | 34 | 25 (10–34) | 26 (13–34) |
| 16 | 24 | 18.5 (8–24) | 19 (10–24) |
| 17 | 26 | 19 (10–26) | 20 (9–26) |

N represents the total number of participants in each one-year age bin. Median *n* (range) represents the median (minimum-maximum) number of participants contributing to valid functional excitation-inhibition ratio (fE/I) paired t-tests across parcels and frequency bands.

**Supplementary Table S2.** EEG-MRI coregistration fits for the pediatric template.

| **Age (years)** | **Mean distance (mm)** |
|---|---|
| 6 | 3.2 |
| 7 | 3.0 |
| 8 | 2.9 |
| 9 | 2.8 |
| 10 | 2.9 |
| 11 | 3.7 |
| 12 | 3.3 |
| 13 | 4.5 |
| 14 | 3.7 |
| 15 | 4.8 |
| 16 | 4.9 |
| 17 | 4.8 |

Values represent the mean distance (mm) between electrode positions and the outer skin surface after rigid coregistration to each pediatric age-specific template.

